# A viral fusogen hijacks the actin cytoskeleton to drive cell-cell fusion

**DOI:** 10.1101/761502

**Authors:** Ka Man Carmen Chan, Sungmin Son, Eva M. Schmid, Daniel A. Fletcher

**Affiliations:** UC Berkeley/UC San Francisco Graduate Group in Bioengineering, Berkeley, CA 94720, USA; Department of Bioengineering & Biophysics Group, University of California, Berkeley, Berkeley, CA 94720, USA; Division of Biological Systems and Engineering, Lawrence Berkeley National Laboratory, Berkeley, CA 94720, USA; Chan Zuckerberg Biohub, San Francisco, CA 94158, USA

## Abstract

Cell-cell fusion, which is essential for tissue development and used by some viruses to form pathological syncytia, is typically driven by fusogenic membrane proteins with tall (>10 nm) ectodomains that undergo conformational changes to bring apposing membranes in close contact prior to fusion. Here we report that a viral fusogen with a short (<2 nm) ectodomain, the reptilian orthoreovirus p14, accomplishes the same task by hijacking the actin cytoskeleton. We show that the cytoplasmic domain of p14 triggers N-WASP-mediated assembly of a branched actin network, directly coupling local force generation with a short membrane-disruptive ectodomain. This work reveals that overcoming energetic barriers to cell-cell fusion does not require conformational changes of tall fusogens but can instead be driven by harnessing the host cytoskeleton.

**Impact Statement:** A viral fusogen drives cell-cell fusion by hijacking the actin machinery to directly couple actin assembly with a short fusogenic ectodomain.

## Introduction

Cell-cell fusion plays a critical role in the development of multicellular organisms, beginning with fertilization and continuing with formation of muscles, osteoclasts, and the placenta in mammals. Viral pathogens, including some members of poxvirus, paramyxovirus, herpesvirus, retrovirus, aquareovirus and orthoreovirus, cause infected cells to fuse with their neighbors, creating syncytia that contribute to disease pathology (Compton & Schwartz, 2017; Domachowske & Rosenberg, 1999; Moss, 2006; Smith, Popa, Chang, Masante, & Dutch, 2009). While the basic steps of membrane fusion have been extensively investigated in the context of enveloped virus entry and SNARE-mediated intracellular vesicle fusion (Sudhof & Rothman, 2009), the molecules and pathways responsible for cell-cell fusion are less well understood. The best studied cell-cell fusogens are those with similarities to enveloped viral fusogens, including syncytin-1 (placental syncytiotrophoblasts formation) (Gong et al., 2005; Renard et al., 2005), Hap2 (conserved in eukaryotic gamete fusion) (Fédry et al., 2017; Feng et al., n.d.; Valansi et al., 2017), and Eff-1 (*C. elegans* epithelial fusion) (Pérez-Vargas et al., 2014; Zeev-Ben-Mordehai, Vasishtan, Siebert, & Grünewald, 2014).

A key feature of viral and cell-cell fusogens is their tall ectodomains, which in their metastable pre-fusion state typically extend more than 10 nm from the membrane. Since the plasma membrane of cells is densely decorated with glycoproteins and glycolipids that could sterically block membranes from getting close enough to fuse, the tall ectodomains of viral and cell-cell fusogens may allow them to reach across the membrane gap and anchor to the apposing membrane, involving insertion of a fusion peptide for Class I viral fusogens or a fusion loop for Class II (Harrison, 2015; Podbilewicz, 2014). Once the fusogen links the two membranes, conformational changes cause the fusogen to fold back, bringing the two membranes into close contact and forming a stable post-fusion structure that promotes membrane fusion (Harrison, 2015; Podbilewicz, 2014; Sapir, Avinoam, Podbilewicz, & Chernomordik, 2008). This conformational change is believed to be sufficient to provide the energy required to overcome the repulsive hydration barrier, which prevents membranes from coming closer than 2 nm (Chernomordik & Kozlov, 2003; Harrison, 2015; Rand & Parsegian, 1989).

However, in other instances of cell-cell fusion, transmembrane proteins required for fusion are short by comparison and do not appear to undergo conformational changes, raising the question of how they bring two plasma membranes into close contact (Figure 1a). One example is the reptilian orthoreovirus fusion protein p14, a non-structural, single-pass transmembrane protein, that is expressed after viral entry. One of seven members of the FAST family of reovirus fusion proteins discovered by Duncan and colleagues (Ciechonska & Duncan, 2014; Corcoran & Duncan, 2004; Dawe & Duncan, 2002; Duncan, Corcoran, Shou, & Stoltz, 2004; Duncan, Murphy, & Mirkovic, 1995; Guo, Sun, Yan, Shao, & Fang, 2013; Kim et al., 2015; Racine et al., 2009; M. Shmulevitz, Epand, Epand, & Duncan, 2004; Maya Shmulevitz & Duncan, 2000; Wilcox & Compans, 1982), p14 has a membrane-disruptive ectodomain that is necessary to drive fusion but extends only 0.7-1.5 nm from the plasma membrane (Corcoran et al., 2006, 2004). This short ectodomain has minimal secondary structure and has no known binding partners that could help to explain how it overcomes the energetic barrier of the crowded plasma membranes and ∼2-nm repulsive hydration barrier to enable fusion (Chernomordik & Kozlov, 2003; Harrison, 2015; Rand & Parsegian, 1989). Yet, expression of p14 alone in cultured cells is sufficient to drive fusion with neighboring naïve cells (Corcoran & Duncan, 2004; Duncan et al., 2004).

**Figure 1.**
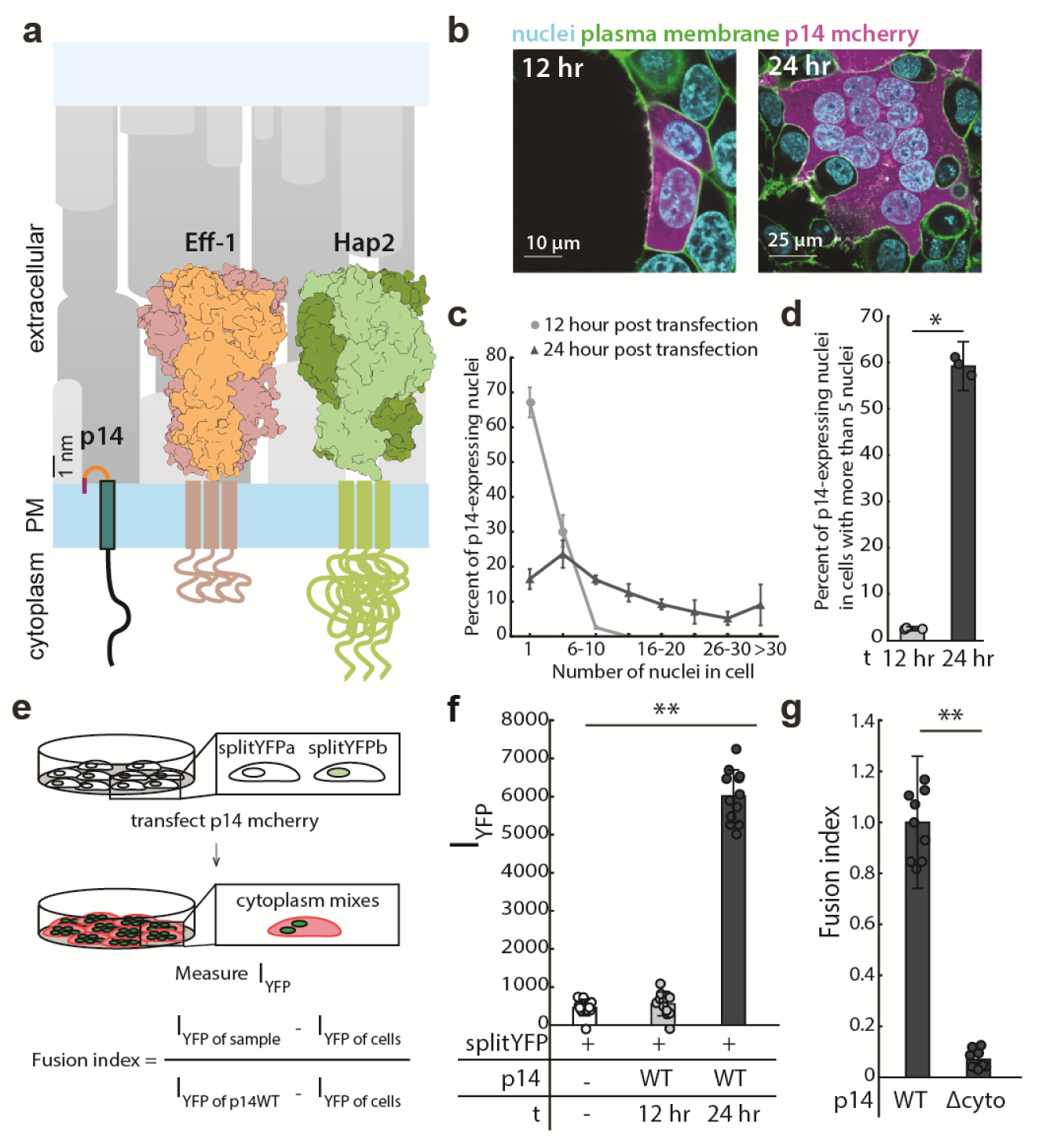
Expression of p14 drives cell-cell fusion and is quantified with splitYFP fluorescent assay. (a) Schematic of fusion-associated small transmembrane protein, p14, in proportion to post-fusion trimeric structure of cell-cell fusogens, Eff-1 (PDB:4OJC) and Hap2 (PDB: 5MF1), on the plasma membrane. (b) Expression of p14 in HEK293T cells drives cell-cell fusion forming large multinucleated cells that increases size and number of nuclei over time. (c) Average nuclei count of multinucleated HEK293T cells expressing p14 at 12 hours and 24 hours with error bars representing standard deviations from 3 independent transfections (See also Figure 1 – source data 1). (d) Percent of p14 expressing nuclei in cells with more than 5 nuclei at 12 hours and 24 hours. p values are two-tailed, two-sample Student’s t-test where *p< 0.01, and error bars represent standard deviations from 3 independent transfections (e) Schematic of splitYFP fluorescence assay to quantify cell-cell fusion. (f) Average YFP fluorescence intensity of HEK293T cells expressing p14 at 12 hours and 24 hours with error bars representing standard deviations from 3 independent transfections of 3 wells each. p values are two-tailed, two-sample Student’s t-test where **p< 0.001 (See also Figure 1 – figure supplement 1c, d). (g) Average fusion index of p14 cytoplasmic truncation mutant with error bars representing standard deviations from 3 independent transfections of 3 wells each. p values are two-tailed, two-sample Student’s t-test where **p< 0.001 (See also Figure 1 – figure supplement 1e, f).

To address the question of how p14 promotes close contact between cells and drives membrane fusion, we studied cell-cell fusion in HEK293T cells transiently expressing p14. We found that the FAST protein, p14 drives cell-cell fusion by hijacking the host cell actin cytoskeleton. Through a phosphorylation-dependent motif in it’s cytoplasmic domain, p14 triggers N-WASP-mediated assembly of a branched actin network, directly coupling local force generation with a short membrane-disruptive ectodomain. This work reveals that overcoming energetic barriers to cell-cell fusion does not require conformational changes of tall fusogens but can instead be driven by harnessing force generated from local actin assembly. This finding points to an alternate means of promoting cell-cell fusion in processes where tall fusogens have not been identified.

## Results

Expression of p14 in HEK293T cells caused the cells to fuse with neighboring wild-type and p14-expressing cells, forming large multinucleated syncytia over the course of 24 hours (Figure 1b, Video 1), like previous reports for other cell types (Corcoran & Duncan, 2004). Partial cleavage of the p14 cytoplasmic tail, which also occurs during reptilian orthoreovirus infection (Top, Barry, Racine, Ellis, & Duncan, 2009), liberates the C-terminus mCherry fluorescent tag from the transmembrane protein and serves as a convenient cytoplasmic marker of p14-expressing cells (Figure 1b, Figure 1 – figure supplement 1a, b, and Video 2). At 12 hours post transfection, 33% of nuclei from p14-expressing cells were in multinucleated cells, with 2% of nuclei in cells with more than 5 nuclei. At 24 hours post transfection, 82% of nuclei from p14-expressing cells were in multinucleated cells, while 59% of nuclei were in cells with more than 5 nuclei (Figure 1c, and d).

To quantify cytoplasmic mixing during p14-mediated cell-cell fusion in a high-throughput manner, we expressed the two halves of splitYFP in two populations of HEK293T cells and mixed the cells together. When fusion occurred, the two halves of splitYFP self-associated in the mixed cytoplasm and fluoresced, allowing quantification by a plate reader (Figure 1e, and Figure 1 – figure supplement 1c and d). Repeating the cell-cell fusion experiments above, the increase in YFP intensity in p14-expressing cells between 12 hours and 24 hours post transfection compared well with the increase in the number of cells with >5 nuclei, as quantified by counting nuclei (Figure 1f).

While the ectodomain of p14 is shorter than typical viral fusogens, its cytoplasmic domain is comparatively long (68 amino acids). To determine how the cytoplasmic domain of p14 might be involved in cell-cell fusion, we first truncated Q70-I125 (p14 Δcyto), retaining a polybasic motif needed for trafficking to the plasma membrane (Parmar, Barry, & Duncan, 2014). Although p14 Δcyto was properly trafficked to the plasma membrane (Figure 1 – figure supplement 1e), cell-cell fusion was abrogated (Figure 1g, and Figure 1 – figure supplement 1f). This is consistent with previous findings (Corcoran & Duncan, 2004), suggesting that p14 may be interacting with cellular components through its cytoplasmic tail to enable cell-cell fusion.

We next investigated whether post-translational modification of the cytoplasmic tail of p14 is for cell-cell fusion. The p14 cytoplasmic tail is mostly disordered but has several tyrosines that could be phosphorylated (Figure 2a, and Figure 2 – figure supplement 1a). To determine if these tyrosines are indeed phosphorylated, we immunoprecipitated p14 and probed with an anti-phosphotyrosine antibody, which confirmed p14 phosphorylation (Figure 2b). Next, we mutated each predicted tyrosine to phenylalanine (Y59F, Y77F, Y96F, Y100F, Y116F) and found that only one mutation (Y116F) decreased cell-cell fusion in our splitYFP assay (Figure 2c and Figure 2 – figure supplement 1b). We then used NetPhos3.1 and Scansite 4.0 to analyze the cytoplasmic tail of p14, and they predicted that Y116 is phosphorylated by c-src kinase (Figure 2d) (Blom, Sicheritz-Pontén, Gupta, Gammeltoft, & Brunak, 2004; Obenauer, 2003). To test this prediction, we mutated all other predicted phosphotyrosines of p14 to phenylalanine (YEY; Y59F, Y77F, Y96F, Y100F) and co-expressed it with constitutively active c-src mutant (CA c-src; Y527F). The experiments showed that Y116 phosphorylation increased, indicating c-src phosphorylates p14 (Figure 2e). Consistent with this, Y116 phosphorylation was also increased when tyrosine phosphatases were broadly inhibited by addition of pervanadate (Figure 2e). Finally, to confirm that c-src is sufficient to phosphorylate Y116, we carried out a modified *in vitro* kinase assay using a peptide including Y116 (P113-N121), along with CA c-src and kinase dead c-src (KD c-src; Y527F/K295R) mutants immunoprecipitated from HEK239T cells (Figure 2 – figure supplement 1c and d). We found that CA c-src was sufficient to phosphorylate p14 cytoplasmic tail peptide, but KD c-src was not (Figure 2f), showing that c-src kinase is necessary and sufficient to phosphorylate Y116 during p14-mediated fusion.

**Figure 2.**
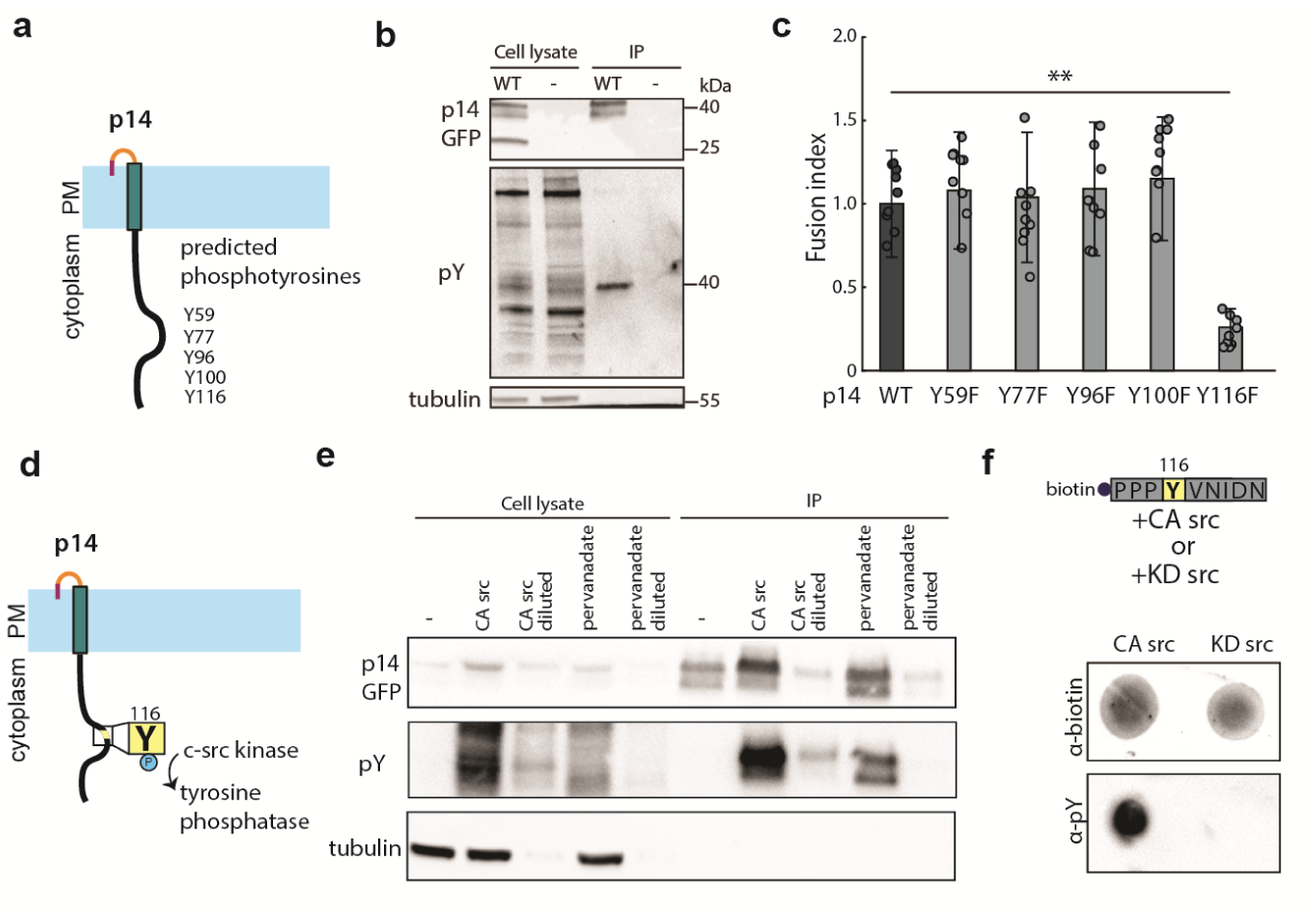
p14 Y116 in the cytoplasmic tail is necessary for cell-cell fusion and is phosphorylated by c-src kinase. (a) Schematic of predicted phosphotyrosines in p14 cytoplasmic tail (See also Figure 2 – figure supplement 1a). (b) Western blot probed with α-phosphotyrosine confirming that p14 WT is phosphorylated. (c) Average fusion index of p14 phosphotyrosine mutants with error bars representing standard deviations from 3 independent transfections of 3 wells each. p values are two-tailed, two-sample Student’s t-test where **p< 0.001 (See also Figure 2 – figure supplement 1b). (d) Schematic of c-src kinase and a tyrosine phosphatase activity on p14 Y116. (e) Western blot probed with α-phosphotyrosine confirming that p14 Y116 phosphorylation is increased with co-expression of constitutively active c-src kinase and with addition of pervanadate. (f) Dot blot of p14 cytoplasmic tail peptide phosphorylated in vitro with constitutively-active (Y527F) and kinase-dead c-src kinase (Y527F, K295R) (See also Figure 2 – figure supplement 1c, d).

To determine which cellular components could be interacting with p14 upon phosphorylation, we used the Eukaryotic Linear Motif (ELM) prediction tool to identify potential binding motifs (Dinkel et al., 2016). ELM predicted that phosphorylated Y116 is bound by the SH2 domain of Grb2 as part of a Grb2 consensus-binding motif, YVNI (Figure 3a). To test this prediction, we carried out a co-immunoprecipitation assay and confirmed that p14 binds to Grb2 (Figure 3b). To determine if Grb2 binding is necessary for p14-mediated cell-cell fusion, we introduced two point mutations that disrupt the predicted Grb2 binding site both individually (Y116F, N118A) and together (FVAI; Y116F/N118A) (Figure 3b). All three mutants severely attenuated cell-cell fusion (Figure 3c, Figure 3 – figure supplement 1a and b), suggesting that Grb2 is important for p14-mediated fusion. To confirm that p14 is sufficient to recruit Grb2, we conjugated biotinylated p14 cytoplasmic tail peptide to streptavidin beads *in vitro* and incubated them with purified Grb2 fluorescently labeled with AF647 (Figure 3 – figure supplement 1c). Consistent with our co-immunoprecipitation results, only phosphorylated Y116 bound to Grb2 (Figure 3d, and Figure 3 – figure supplement 1d). When p14 Y116 phosphorylation is increased in cells with either co-expression of CA c-src or addition of the phosphatase inhibitor pervanadate, GFP-labeled Grb2 co-localized with p14 at the plasma membrane (Figure 3e). However, Grb2 did not co-localize with p14 at the plasma membrane when the Grb2 binding site was mutated to FVAI and co-expressed with CA src or treated with pervanadate (Figure 3e, and Figure 3 – figure supplement 1e, f, g).

**Figure 3.**
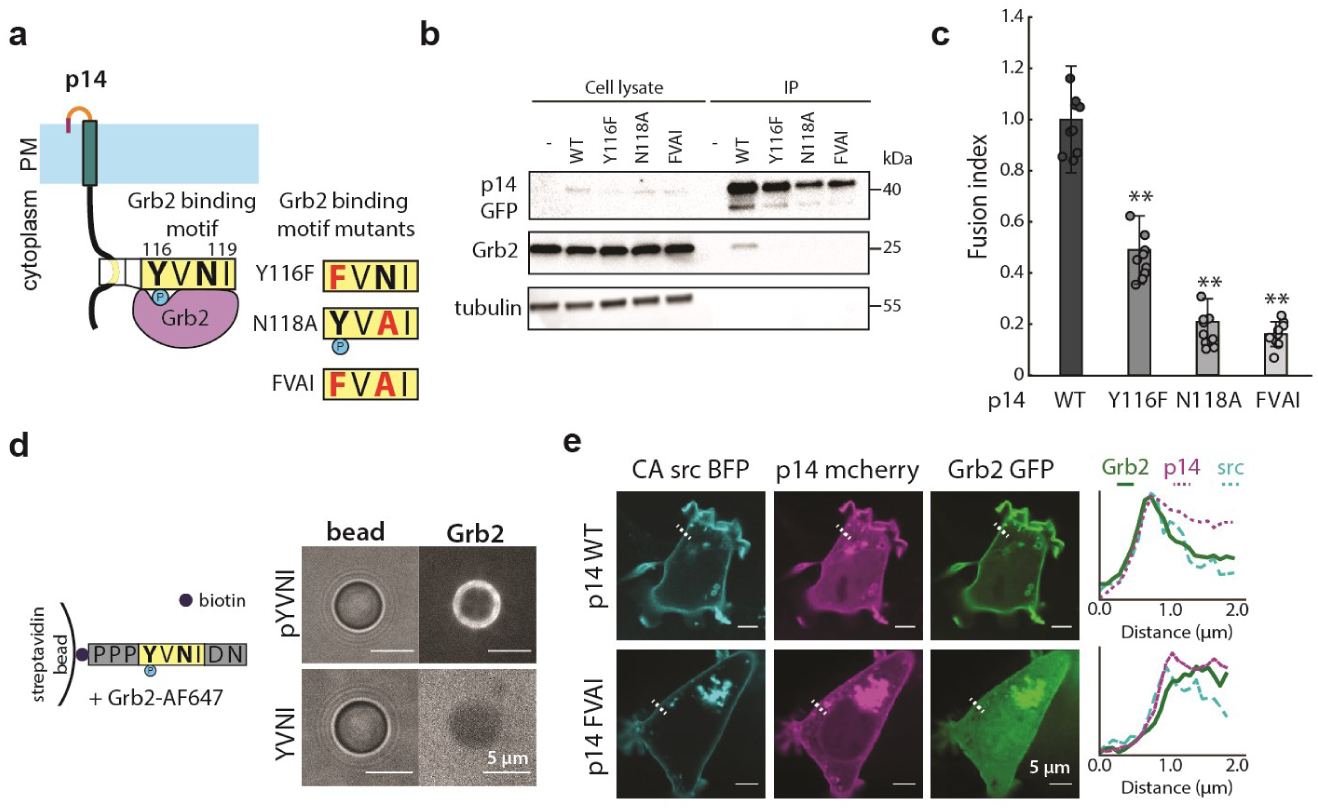
p14 Y116 in the cytoplasmic tail binds to Grb2. (a) Schematic of p14 mutants that disrupt predicted Grb2 binding motif. (b)Western blot of co-immunoprecipitation of p14 with Grb2 (lane 7) and p14 mutants, Y116F, N118A, FVAI, that does not bind Grb2 (lane 8, 9, 10). (c) Average fusion index of p14 mutants with error bars representing standard deviations from 3 independent transfections of 3 wells each. p values are two-tailed, two-sample Student’s t-test where **p< 0.001 (See also Figure 3 – figure supplement 1a, b). (d) Streptavidin beads with biotinylated phosphorylated and non-phosphorylated Y116 p14 cytoplasmic tail peptide encoding (P113-N121) binds and did not bind to purified Grb2 respectively (See also Figure 3 – figure supplement 1c, d). (e) Confocal images of Grb2 enrichment to the plasma membrane of cells co-expressing p14 WT with constitutively active c-src kinase. A line scan of fluorescence intensity of each protein along the indicated white line (See also Figure 3-figure supplement 1e, f, g).

Having shown that Grb2 binds to the p14 cytoplasmic tail in a phosphorylation-dependent manner, we next sought to determine mechanistically how Grb2, an adaptor protein with two SH3 domains, plays a role in p14-mediated cell-cell fusion. The N-terminal SH3 domain of Grb2 binds to SOS, activating Ras, which in turn activates Raf kinase and the MAPK-ERK1/2 pathway, while the C-terminal SH3 domain of Grb2 binds to the actin nucleation promoting factor N-WASP, which binds to Arp2/3 and nucleates branched actin assembly (Figure 4a). To determine if one or both pathways are important for fusion, we first treated cells expressing p14 with sorafenib tosylate, an inhibitor of Raf kinase, but found no effect on cell-cell fusion at up to 100 times the IC50 (Figure 4b). We next considered whether branched actin networks could be directly involved in p14-mediated cell-cell fusion. Building on previous work showing that cytochalasin D disrupts fusion of p14-expressing cells (Salsman, Top, Barry, & Duncan, 2008), we treated p14-expressing cells with wiskostatin, an inhibitor of N-WASP, and found that fusion was significantly reduced (Figure 4b). We then treated p14-expressing cells with CK-666 to inhibit the Arp2/3 complex, which forms the branches in branched actin networks, and found that fusion was reduced in a dose-dependent manner (Figure 4b). In contrast, treating p14-expressing cells with smifH2, an inhibitor of formins, enhanced cell-cell fusion (Figure 4b), perhaps due to increased branched actin assembly that has been observed when formins are broadly inhibited (Burke et al., 2014).

**Figure 4.**
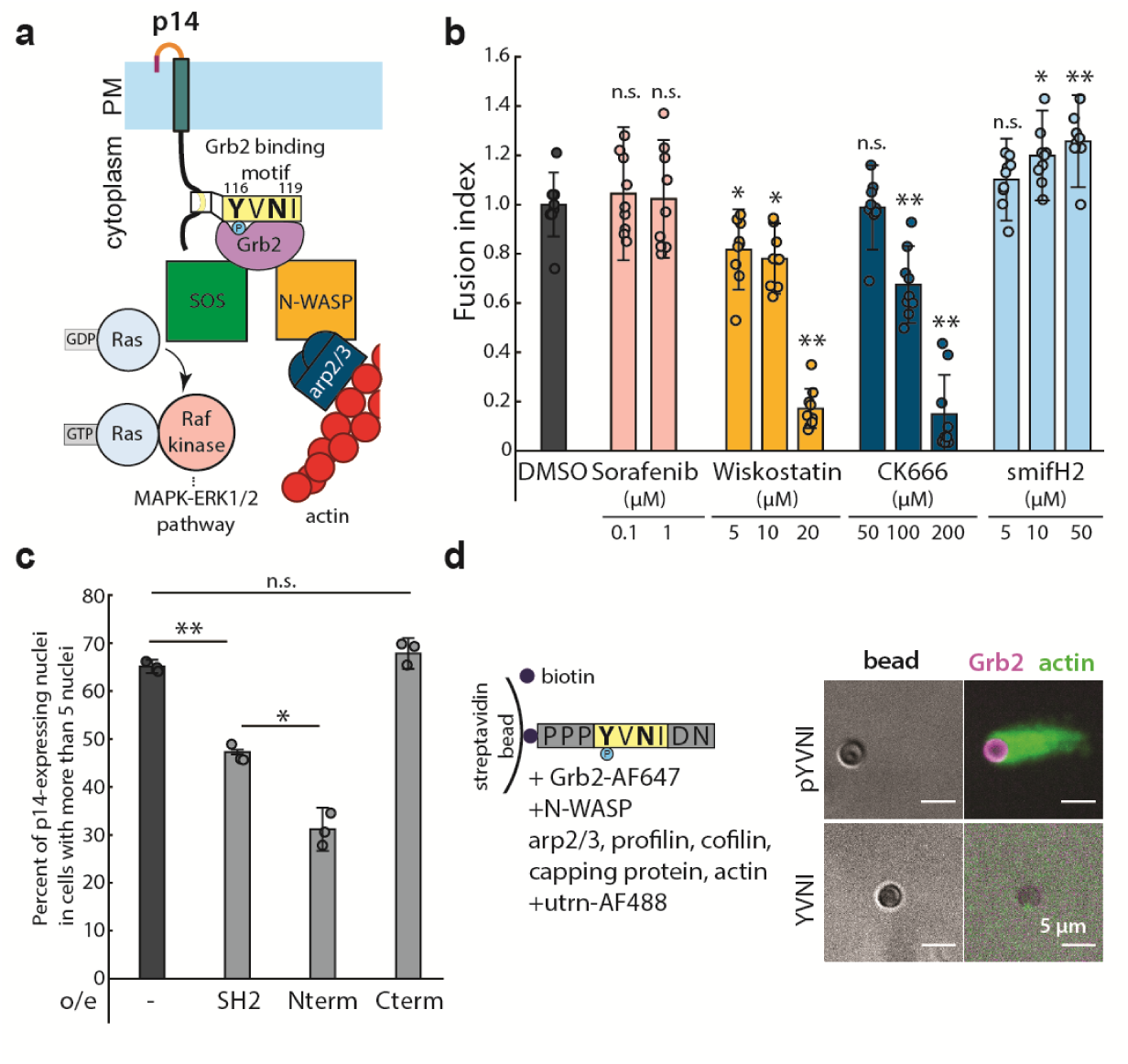
N-WASP-dependent assembly of branched actin network is necessary for cell-cell fusion. (a) Schematic of Grb2 binding to two potential downstream effectors, SOS and N-WASP (b) Extent of cell-cell fusion quantified with splitYFP fluorescence assay of p14 expressing cells treated sorafenib tosylate targeting Raf kinase, wiskostatin targeting N-WASP, CK666 targeting Arp2/3 and smifH2 targeting formins, normalized to that of p14 WT treated with vehicle control, DMSO. Error bars indicate standard deviations from 3 independent transfections of 3 wells each. p values are two-tailed, two-sample Student’s t-test to DMSO where *p< 0.01, and **p<0.001. (c) Average percent of p14 expressing nuclei in cells with more than 5 nuclei of p14-expressing HEK293T WT cells and HEK293T cells overexpressing Grb2 SH2 domain, N-terminus SH2-SH3 mutant and C-terminus SH2-SH3 mutant. p values are two-tailed, two-sample Student’s t-test where *p< 0.01, and **p<0.001. Error bars represent standard deviations from 3 independent transfections (See also Figure 4 – figure supplement 1a, b, c, d, and Figure 4 – source data 1). (d) In vitro actin bead motility of phosphorylated p14 cytoplasmic tail peptide conjugated to streptavidin beads in a purified actin motility mixture supplemented with Grb2. Polymerized actin is visualized with AlexaFluor488-labeled utrophin actin binding domain (See also Figure 4 – figure supplement 1e).

To test whether the reduction in cell-cell fusion with wiskostatin and CK-666 was the result of a direct link between N-WASP and p14 or a more general inhibition of actin activity, we over-expressed Grb2 mutants that can only bind to either SOS or N-WASP by truncating either the N- or C-terminal SH3 domains. We found that both of these Grb2 mutants bound to p14 WT and co-localized with phosphorylated p14 in pervanadate-treated live cell images (Figure 4 – figure supplement 1a), confirming that the mutations did not disrupt interactions with p14. However, when the Grb2 N-terminal mutant that can bind only to SOS (Nterm) was over-expressed, the extent of p14-mediated cell-cell fusion was reduced, similar to when endogenous Grb2 binding was reduced by overexpression of the Grb2 SH2 domain (Figure 4c, Figure 4 – figure supplement 1b, c and d). When a Grb2 C-terminal mutant that can bind only to N-WASP (Cterm) was over-expressed, p14-mediated cell-cell fusion was restored to a level comparable to that of endogenous Grb2 in WT cells (Figure 4c, Figure 4 – figure supplement 1b, c, and d).

To determine whether N-WASP binding to Grb2 and the p14 cytoplasmic tail is sufficient to nucleate actin assembly, we used an *in vitro* actin-based motility assay. We bound biotinylated p14 cytoplasmic tail peptides to streptavidin beads in a purified actin motility mixture containing N-WASP (lacking the EVH1 domain), Arp2/3, profilin, cofilin, capping protein and actin, supplemented with Grb2 (Figure 4 – figure supplement 1e). When Y116 of the p14 cytoplasmic tail peptide was phosphorylated, actin tails were nucleated from the bead (Figure 4d), but when Y116 was not phosphorylated, actin tails were not observed (Figure 4d). This confirms that Grb2 is necessary and sufficient to recruit N-WASP to the p14 cytoplasmic tail and can nucleate localized branched actin networks when p14 is present.

We next investigated whether branched actin network assembly must be directly coupled to the fusogenic ectodomain, or whether the fusogenic ectodomain can simply be present in the same membrane as actin assembly by the cytoplasmic tail of p14. To test the necessity for direct coupling, we co-expressed the ectodomain deletion mutant (Δecto; ΔM1-T35), which traffics to the plasma membrane and binds Grb2 (Figure 5 – figure supplement 1a and b) and the cytoplasmic tail deletion mutant (Δcyto) in the same cell. Interestingly, we found that cell-cell fusion was abolished (Figure 5a, and Figure 5 – figure supplement 1c), despite the presence of both halves of p14. This indicates that localized actin assembly is necessary for p14-mediated fusion and suggests a reason why native Arp2/3-generated branched actin networks formed at cell-cell contacts are not sufficient to cause spontaneous cell-cell fusion.

**Figure 5.**
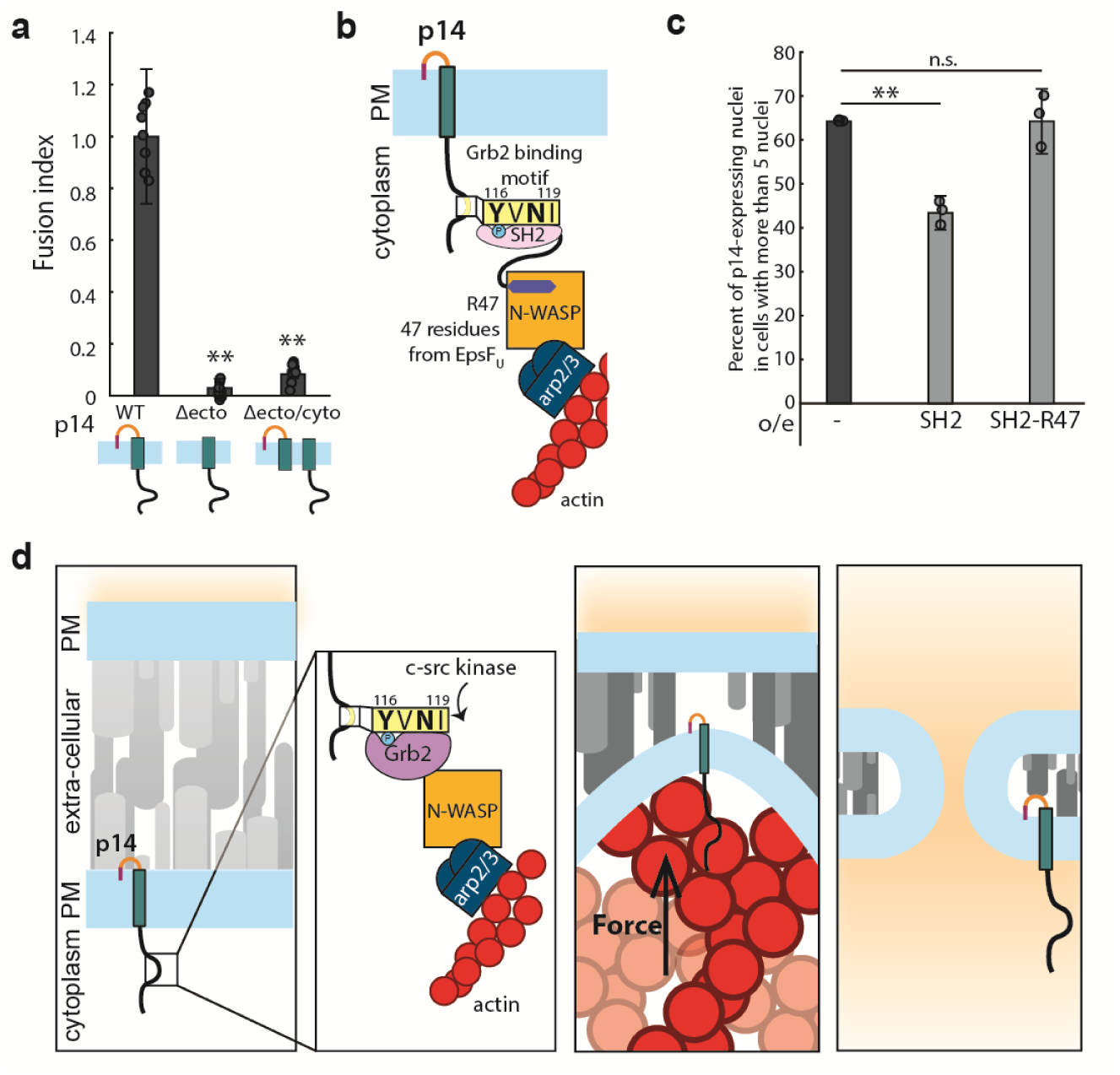
Branched actin assembly directly coupled to p14 cytoplasmic tail drives cell-cell fusion. (a) Extent of cell-cell fusion quantified with splitYFP fluorescence assay of p14 truncation mutants normalized to that of p14 WT. p values are two-tailed, two-sample Student’s t-test to p14 WT where *p< 0.01 and **p<0.001. Error bars indicate standard deviations from 3 independent transfections of 3 wells each (See also Figure 5 – figure supplement 1a, b, c). (b) Schematic of fusion protein coupling actin assembly to p14 cytoplasmic tail consisting of Grb2 SH2 domain and 47 residues from EpsFU. (c) Average percent of p14 expressing nuclei in cells with more than 5 nuclei of p14-expressing HEK293T WT cells and HEK293T cells overexpressing Grb2 SH2 domain and SH2-R47. p values are two-tailed, two-sample Student’s t-test where **p<0.001. Error bars represent standard deviations from 3 independent transfections (See also Figure 5 – figure supplement 1d, e, f, g, and Figure 5 – source data 1). (d) Proposed mechanism of coupling of force generation of actin assembly at p14 cytoplasmic tail with the small ectodomain to drive cell-cell fusion.

If a localized pushing force is necessary and sufficient for the fusogenic activity of p14, then it should be possible to drive cell-cell fusion by creating an alternate link between p14 and branched actin network assembly. To test this idea, we engineered a fusion protein that binds to p14 consisting of Grb2 SH2 domain and a 47-residue peptide from EpsFU of enterohemorrhagic *E. coli* (EHEC) (Figure 5b). This 47-residue peptide binds to and relieves the auto-inhibition of endogenous N-WASP, nucleating branched actin network (Cheng, Skehan, Campellone, Leong, & Rosen, 2008; Sallee et al., 2008). Fusion of this peptide with the Grb2 SH2 domain, which we confirmed binds to phosphorylated p14 (Figure 5 – figure supplement 1d), enables binding to WT p14. When we expressed this fusion protein (SH2-R47) together with p14 in HEK293T, cell-cell fusion was significantly higher than when the SH2 domain lacking R47 was expressed to a similar level (Figure 5c, Figure 5 – figure supplement 1e, f and g). This result demonstrates that direct coupling of branched actin assembly to p14 is necessary and sufficient for fusion.

## Discussion

Taken together, these experiments reveal a viral pathogen that hijacks branched actin network assembly to drive cell-cell fusion, reminiscent of how the pathogens *Listeria monocytogenes* and vaccinia virus hijack branched actin network assembly to move within and between cells (Welch & Way, 2013). Here we show that the reptilian orthoreovirus fusogen p14 accomplishes this by presenting a c-src kinase substrate in its cytoplasmic tail, binding the host cell adaptor protein Grb2, and nucleating branched actin assembly through the host nucleation promoting factor N-WASP. Since the p14 ectodomain extends only 0.7-1.5 nm from the plasma membrane (Corcoran et al., 2006), it cannot interact with apposing membrane through the ∼2 nm repulsive hydration barrier that prevents two membranes from coming together. We propose that the primary role of this branched actin network assembly is to physically push the p14 ectodomain into close contact with a neighboring cell (Figure 5d), a step carried out by conformational changes of tall fusogens in previously studied examples of plasma and viral membrane fusion. Localized pushing by p14’s cytoplasmic domain enables direct interaction between the apposing membrane and p14’s membrane-disruptive ectodomain, which contains hydrophobic residues and myristoylation that are known to be necessary for fusion (Corcoran & Duncan, 2004; Top et al., 2009). The actin cytoskeleton may also play other roles, such as clustering p14 ectodomains (Köster & Mayor, 2016), changing local membrane curvature and tension (Kozlov & Chernomordik, 2015), blocking fusion pore closure and expanding the fusion pore (Kozlov & Chernomordik, 2015), that could further promote membrane fusion.

The actin cytoskeleton and its accessory proteins have been implicated in various cell-cell fusion events, including formation of signaling scaffolds, protrusive structures, and mechanical resistance in *C. elegans* epithelial fusion (Yang et al., 2017), osteoclast fusion (Oikawa et al., 2012), and myoblast fusion (Chuang et al., 2019; Kim et al., 2015; Shilagardi et al., 2013). Here we demonstrate an additional role for the actin cytoskeleton in cell-cell fusion – physically forcing a short fusogenic ectodomain through dense cell surface proteins and into contact with an apposing membrane. This force-mediated fusion mechanism could be relevant for other instances of cell-cell fusion involving short transmembrane proteins, such as myomixer, a short extracellular peptide, which is required for myoblast fusion and likely extend only a few nanometers from the plasma membrane (Bi et al., 2018; Leikina et al., 2018; Millay et al., 2013; Quinn et al., 2017; Sampath, Sampath, & Millay, 2018; Zhang et al., 2017). Forces generated by the cytoskeleton, rather than by conformational changes in a tall fusogen, may be used to bring two membranes into close contact during myoblast fusion, as well as in other cell-cell fusion events such as macrophage giant cell formation, where no fusogens have been identified yet.

## Acknowledgments

The authors would like to thank R. Duncan and E. Chen for helpful discussion and the Fletcher Lab members, including M.H. Bakalar, B.D. Belardi, A.R. Harris, and M.D. Vahey, for useful feedback and technical consultation.

## Contributions

Conceptualization, K.M.C.C., S.S., E.M.S., and D.A.F.; Methodology, K.M.C.C. and S.S.; Validation, K.M.C.C.; Formal analysis, K.M.C.C.; Investigation, K.M.C.C. and S.S.; Resources, K.M.C.C.; Writing – original draft preparation, K.M.C.C. and D.A.F.; Writing – review & editing, K.M.C.C., S.S., E.M.S., and D.A.F.; Visualization, K.M.C.C.; Supervision, D.A.F.; Project administration, D.A.F.; Funding acquisition, S.S., E.M.S., and D.A.F..

## Data and materials availability

All data generated or analyzed during this study are included in the manuscript and supporting files.

## Materials and methods

### Cloning

Reptilian reovirus membrane fusion protein, p14 (Accession number: Q80FJ1), was synthesized and inserted into mammalian expression vector pcDNA3.1 with C-terminus tags (mcherry, eGFP). Point mutations and truncations were introduced with primers.

splitYFPa and splitYFPb were amplified from pBiFC-bJun-YN155 and pBiFC-bFos-YC155 (a kind gift from Tom Kerppola) and inserted into lentiviral transfer plasmid, pHR, with Gibson assembly.

Constitutively active chick-src kinase was amplified from pLNCX chick src Y527F and inserted into pcDNA 3.1 with linker (GGGS) and C-terminus tags (FLAG and mTagBFP2). pLNCX chick src Y527F was a gift from Joan Brugge (Addgene plasmid # 13660). K295R was introduced to constitutive active chick-src kinase to render it kinase dead with primers.

cDNA of Human Grb2 (GE Dharmacon, cloneID: 3345524) was amplified and inserted into pGEX4T2 with a N-terminus GST tag and TEV cleavage site for purification of Grb2. For overexpression of Grb2 mutants, IRES Puromycin was amplified from pLV-EF1a-IRES-Puro (a gift from Tobias Meyer, Addgene plasmid #85132) and inserted into lentiviral pHR backbone to create pHR-IRES-Puro. Grb2 N-terminus SH3 domain and SH2 domain (N-termSH3, 1-159), Grb2 C-terminus SH3 domain and SH2 domain (C-termSH3, 58-217), and Grb2 SH2 domain (58-159) were amplified from cDNA of Human Grb2 and inserted into pHR-IRES-Puro with C terminus FLAG tag.

47 residues from EpsF(U) of enterohemorrhagic *E. coli* (268–314) was synthesized and inserted with GGGS linker downstream of Grb2 SH2 domain (58-159) and FLAG tag into pHR-IRES-Puro.

### Cell culture, transfection and generation of mutant Grb2 overexpression cells

HEK293T cells were obtained from UCSF Cell Culture Facility. HEK293T cells were grown in DMEM (Life Technologies) supplemented with 10% heat-inactivated FBS (Life Technologies) and 1% Pen-Strep (Life Technologies), at 37 °C, 5% CO_2_. Cells were negative for mycoplasma as verified with Mycoalert mycoplasma detection kit (Lonza).

Cells were transfected with TransIT-293 (Mirus Bio) according to manufacturer’s instructions.

To over-express Grb2 mutants to compete with endogenously expressed Grb2 and SH2 actin nucleators, pHR-Grb2NtermSH3-FLAG-IRES-Puro, pHR-Grb2CtermSH3-FLAG-IRES-Puro, pHR-Grb2SH2-FLAG-IRES-Puro, pHR-SH2-FLAG-R47 were co-transfected with second generation packaging plasmids, pMD2.G and p8.91 in HEK293T to generate lentivirus.

HEK293T cells were transduced with lentivirus, and 24 hours post transduction selected with 3 μg/ml puromycin (Clontech) to select for mutant Grb2 expression. Cultures were maintained in 3 μg/ml puromycin.

### splitYFP cell-cell fusion assay

pHR-splitYFPa and pHR-splitYFPb were co-transfected with second generation packaging plasmids, pMD2.G and p8.91 in HEK293T to generate lentivirus. WT HEK293T cells were transduced with splitYFPa and splitYFPb lentivirus. The cells were passaged for at least a week before use in cell-cell fusion assay.

To quantify cell-cell fusion, HEK293T cells stably expressing splitYFPa and splitYFPb were mixed at 50:50 ratios and 1.33×10^5^ of cells were plated into each well of 48 well plate. The next day, the cells were transfected with TransitIT-293 (Mirus Bio). 18 hours post transfection, cells were moved to 30°C, 5% CO_2_ incubator to mature the splitYFP fluorophore. 24 hours post transfection, cells were lifted with 150 μl of 2mM EDTA and placed into 96 well black bottom plate. splitYFP was excited at 510 nm and emission at 530 nm was quantified using a plate reader (Tecan).

Fusion index was quantified as (I_sample – I_cell) /(I_p14WT – I cell), where I_cell is the YFP intensity of non-transfected HEK293T cells expressing splitYFPa and splitYFPb, I_sample is the YFP intensity of HEK293T cells transfected with plasmid as specified, I_ p14WT is the YFP intensity of HEK293T cells transfected with p14 WT and treated with DMSO as vehicle control. Average and standard deviation of fusion index is calculated from 3 independent transfections of 3 wells each. Statistical significance was determined using two-tailed, two-sample Student’s t-test.

### Nuclei count

3.8×10^5^ cells were plate into 24 well plates and transfected the next day with designated plasmid with TransIT-293 (Mirus Bio) according to manufacturer’s instructions. 2 hours post transfection, cells were lifted with 150 μl of 2 mM EDTA, re-suspended with 850 μl of media, and 300 μl of cell suspension was plated onto a fibronectin-coated glass bottom chamber (Cell-vis). At 6 hours and 18 hours post transfection, cells were transferred to 30°C, 5% CO_2_ incubator. After 6 hours incubation at 30°C, nuclei were labeled with 0.6% Hoescht 33342 (Life Technologies), and plasma membrane were labeled with 0.05% CellMask Deep Red (Thermo Fisher Scientific) for 20 min at 37°C. Cells were imaged using spinning disk confocal microscopy. About 80-100 random field of views are taken for each sample to image almost the entire imaging well, and the number of nuclei in p14 expressing cells are manually counted. Average and standard deviation of binned nuclei count is calculated from 3 independent transfections. Statistical significance was determined using two-tailed, two-sample Student’s t-test.

### Drug treatment

To broadly inhibit the protein-tyrosine phosphatases, pervanadate is prepared by incubating 10 mM sodium orthovanadate with 0.15% hydrogen peroxide in 20 mM HEPES for 5 min at room temperature. Pervanadate is neutralized with catalase and added to cells immediately. For western blot, cells were lysed 10 mins after pervanadate addition, for live-imaging, cells were imaged immediately after pervanadate addition.

To perturb the actin cytoskeleton, 4 hours post transfection, the media was replaced with complete media supplemented with cytoskeletal drugs CK-666 (Sigma Aldrich), Wiskostatin (Krackeler Scientific) and smifH2 (EMD Millipore) at specified concentrations. DMSO was used as vehicle control. 18 hours post transfection, splitYFP was matured at 30°C. At 24 hours post transfection splitYFP fluorescence was quantified as described above.

To inhibit Raf kinase, 4 hours post transfection, the media was replaced with compete media supplemented with sorafenib tosylate (Selleckchem). DMSO was used as a vehicle control. 18 hours post transfection, splitYFP was matured at 30°C. At 24 hours post transfection splitYFP fluorescence was quantified as described above.

### Co-immunoprecipitation

HEK293T were transfected with specified plasmids. 17-24 hours post transfection, HEK293T cells were washed with 1 mM CaCl_2_/PBS, lifted off the dish with 2 mM EDTA/PBS, pelleted and lysed by incubating in lysis buffer (150 mM NaCl, 25 mM HEPES, 1 mM EDTA, 0.5% NP-40, 1x PhosSTOP phosphatase inhibitor (Roche), 1x HALT protease inhibitor (Thermo Fisher Scientific) for 30 min, and bath sonicated in ice for 3 min. Cell debris was pelleted at 18,000 rcf for 10 min. Cell lysate were precleared with 15 μl of GFP-Trap (Chromotek) for 30 min at 4°C, and incubated with 15 μl of fresh GFP-Trap beads overnight at 4°C. The beads were washed with lysis buffer five times, before boiled in Laemmli sample buffer and separated on 4-20% acrylamide gradient gels by SDS-PAGE. Proteins were transferred onto nitrocellulose membrane and probed with primary antibodies, α-Grb2 (1:5000, Clone 81/Grb2, BD Biosciences), α-tubulin (1:5000, Clone YL1/2, Thermo), α-pTyr (1:2000, Phospho-Tyrosine (P-Tyr-1000) MultiMab™Rabbit mAb mix #8954, Cell Signaling Technology), α-GFP (1:10000, Clone 3E6, Life Technologies or 1:5000, A21312, Life Technologies), and secondary antibodies, α-mouse HRP (1:10,000, Upstate Biotechnology or 1:5000, Jackson Labs), α-rabbit HRP (1:5000, 65-6120, Thermo Fisher), α-rat AlexaFluor 647(1:5000, Life Technologies). Western blots were imaged on ChemiDoc (Bio-Rad).

### Membrane fractionation

HEK293T were transfected with p14 WT, p14 Y116F/N118A, and p14 Δcyto. 18 hours post transfection, the cells were washed with 1 mM CaCl_2_/PBS, lifted off the dish with 2 mM EDTA/PBS. Cells were pelleted at 200 rcf for 5 min and re-suspended in fractionation buffer (20 mM HEPES, 10 mM KCl, MgCl_2_, 1 mM EDTA, 1 mM EGTA, 1 mM TCEP, 1x HALT protease inhibitor(Thermo Fisher Scientific)). The cell suspension lysed with five freeze/thaw cycles. Nuclei were pelleted via centrifugation (700 rcf, 5 min), and mitochondria were pelleted at 10,000 rcf, 5 min. The supernatant was then centrifuged at 100,000 rcf for an hour at 4°C to separate the membrane and cytoplasmic fraction. The membrane pellet was washed once in fractionation buffer and re-centrifuged at 100,000 rcf for an hour. The cell lysate, cytoplasmic fraction, and membrane pellet was boiled in Laemmli sample buffer, and separated on 4-20% acrylamide gradient gels by SDS-PAGE. Proteins were transferred onto nitrocellulose membrane and probed with primary antibodies, α-tubulin (1:5000, Clone YL1/2, Thermo Fisher Scientific), α-GFP (1:5000, A-21312, Life Technologies), and secondary antibodies, α-rabbit HRP (1:5000, 65-6120, Thermo Fisher) and α-rat AlexaFluor 647(1:5000, Life Technologies). Western blots were imaged on a ChemiDoc (Bio-Rad).

### Protein purification

GST-TEV-Grb2 (human) was expressed and purified from *E. coli* as previously described (Su et al., 2016).

N-WASP (ΔEVH1) was a kind gift from D. Wong and J. Taunton (University of California, San Francisco). Actin was purified from rabbit skeletal muscle as previously described (Spudich & Watt, 1971). Capping protein was a kind gift from S. Hansen and D. Mullins (University of California, San Francisco). Arp2/3 was purchased from Cytoskeleton, Inc. Profilin, cofilin, and utrophin actin binding domain (1-261) were purified as previously described(Bieling et al., 2016).

### Motility assay

Similar to a previously described motility assay(Okrut, Prakash, Wu, Kelly, & Taunton, 2015), 2 μl of 0.5% 3 μm streptavidin polystyrene beads (Bangs Laboratories) are incubated with 1 μM biotin-p14 cytoplasmic tail peptide in 10 mM HEPES (pH 7.5), 1 mg/ml BSA and 50 mM KCl for 10 min at room temperature. Peptide-coated beads are diluted eight-fold into motility buffer (10 mM HEPES, 2 mM MgCl2, 50 mM KCl, 50 mM NaCl, 1 mg/ml BSA, 2.5 mM ATP, 5 mM TCEP), containing 0.1 μM Grb2 (20% labeled), 0.2 μM N-WASP, 9 μM actin, 0.075 μM arp2/3, 0.05 μM capping protein, 2.6 μM profilin, 3.5 μM cofilin and incubated for 15 min at room temperature while rotating. 300 nM utrophin-AF488 is added to the mixture, and incubated for 5 min, before imaging.

### In vitro kinase assay

As previously described (Dagliyan et al., 2016), HEK293T is transiently transfected with constitutively active chick src (Y527F) kinase and kinase dead (Y527F/ K295R) src kinase with C-terminus FLAG tag with TransIT-293 (Mirus). 24 hours post transfection, cells were washed with 1 mM CaCl_2_/PBS, and lifted with 2 mM EDTA. Cells were pelleted, and lysed in 20 mM HEPES-KOH, 50 mM KCl, 100 mM NaCl, 1 mM EGTA, 1% NP-40, 1x PhosSTOP phosphatase inhibitor (Roche) and 1x HALT protease inhibitor (Thermo Scientific) for 30 min, 4°C while rotating. Cell debris was pelleted at 3000 g, 10 min, and FLAG-tagged kinase were immunoprecipitated with 3 μg of α-FLAG (M2 clone, Sigma) and 50 μl of Protein-G Dynabeads (Thermo Scientific) for 2 hours, 4°C, while rotating. Beads were washed twice with intracellular buffer (20 mM HEPES-KOH, 50 mM KCl, 100 mM NaCl, 1 mM EDTA, 1% NP-40), and twice with kinase buffer (25 mM HEPES, 5 mM MgCl_2_, 5 mM MnCl_2_, 0.5 mM EGTA). Beads were re-suspended in kinase buffer, supplemented with 0.2 mM ATP and 1.5 mM biotin-p14 cytoplasmic tail peptide, and incubated for 1 hour at room temperature. Protein-G dynabeads were removed, and the supernatant is incubated with 10 μl streptavidin magnetic beads (Pierce) for 30 min, room temperature. Streptavidin magnetic beads were washed twice with kinase buffer, and boiled in sample buffer. Sample are dotted onto nitrocellulose membrane (Bio-Rad), and blocked with 5% BSA, and probed with α-pTyr (1:5000, Phospho-Tyrosine (P-Tyr-1000) MultiMab™Rabbit mAb mix #8954, Cell Signaling Technology) and α-biotin-AF647 (1:5000, BK-1/39, Santa Cruz Biotechnology) in 5% BSA overnight at 4°C. Blots were washed 3 times, 5 min each with TBST, and probed with secondary antibody, α-rabbit HRP (1:5000, 65-6120, Thermo Fisher) and washed 3 times, 15 min each. Blots were imaged on Chemi-Doc (Bio-rad).

### Imaging

All live cells were maintained at 37°C, 5% CO_2_ with a stage top incubator (okolab) during imaging.

For confocal microscopy, cells were imaged with a spinning disk confocal microscope (Eclipse Ti, Nikon) with a spinning disk (Yokogawa CSU-X, Andor), CMOS camera (Zyla, Andor), and either a 4x objective (Plano Apo, 0.2NA, Nikon) or a 60x objective (Apo TIRF, 1.49NA, oil, Nikon). For total internal reflection fluorescence (TIRF) microscopy, cells were imaged with TIRF microscope (Eclipse Ti, Nikon), 60x objective (Apo TIRF, 1.49NA, oil, Nikon) and EMCCD camera (iXON Ultra, Andor). Both microscopes were controlled with Micro-Manager. Images were analyzed and prepared using ImageJ (National Institutes of Health). To capture large multinucleated cells, such as in Fig. S1., multiple fields of view were stitched together using Grid/Collection Stitching plugin in Fiji.

## Figure Supplements

**Figure 1 – figure supplement 1.**
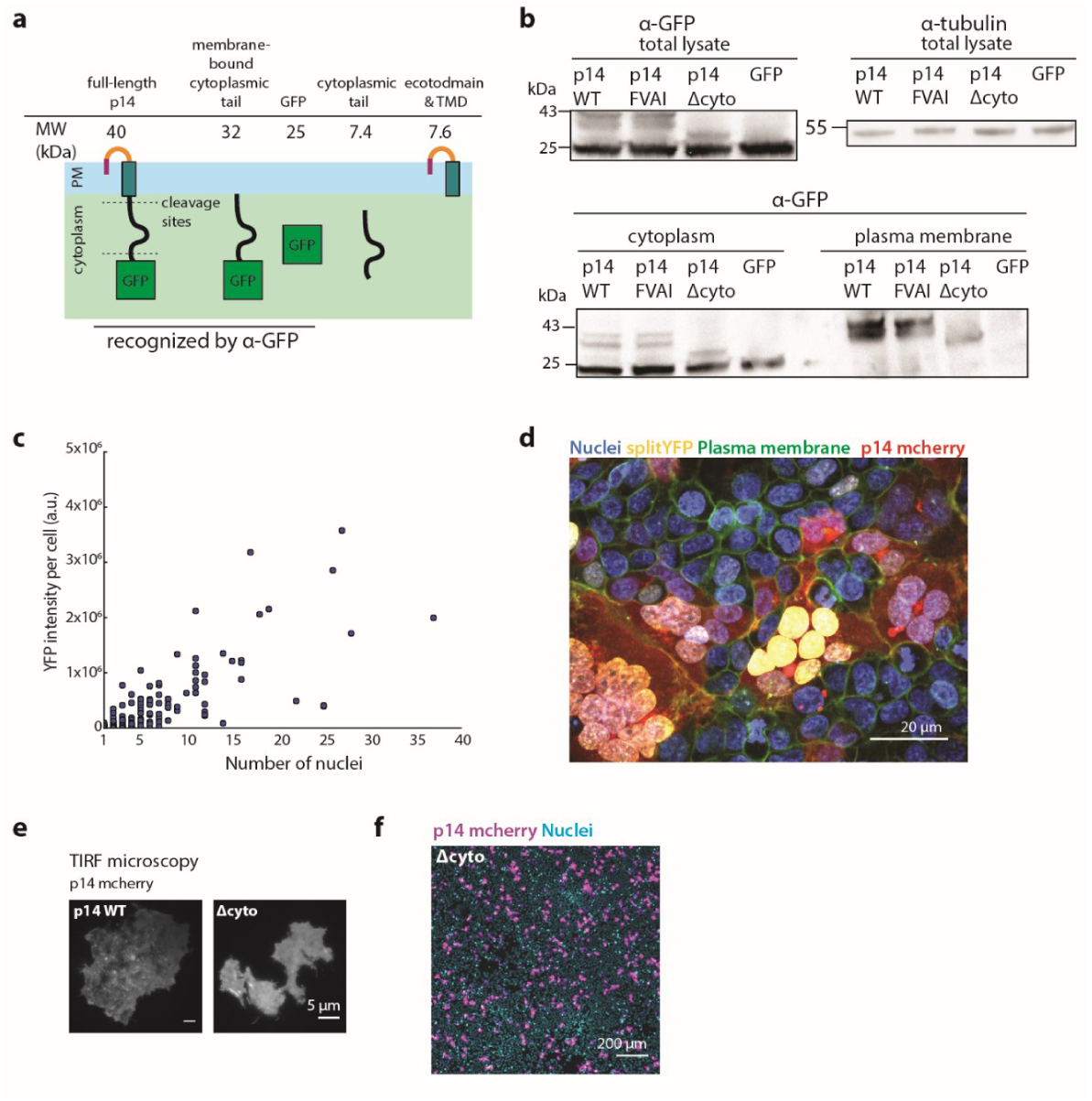
Characterization of p14 cytoplasmic tail cleavage and splitYFP cell-cell fusion assay. (a) Schematic of hypothesized cleavage sites in p14 cytoplasmic tail based on molecular weights of cleaved fragments. (b) Membrane fractionation of p14 WT, p14 FVAI, Δcyto and GFP only, showing similar expression of each construct at the plasma membrane and molecular weights cleaved fragments. (c) YFP intensity per cell vs number of nuclei in cell. Each point represents a single cell. (d) Representative confocal image of splitYFP cells expressing p14 WT mcherry. Nuclei are labeled with Hoechst 33342, plasma membrane are labeled with CellMaskDeepRed. splitYFP has FOS and JUN coiled-coiled motif that directs splitYFP to the nucleus. (e) p14 WT and p14 Δcyto labeled with C-terminus mcherry tag are trafficked to the plasma membrane as visualized with TIRF microscopy. (f) Representative field of view of HEK293T cells expressing p14 WT and p14 Δcyto with p14 labeled with C-terminus mcherry (magenta), nuclei stained with Hoechst 33342 (cyan).

**Figure 2 – figure supplement 1.**
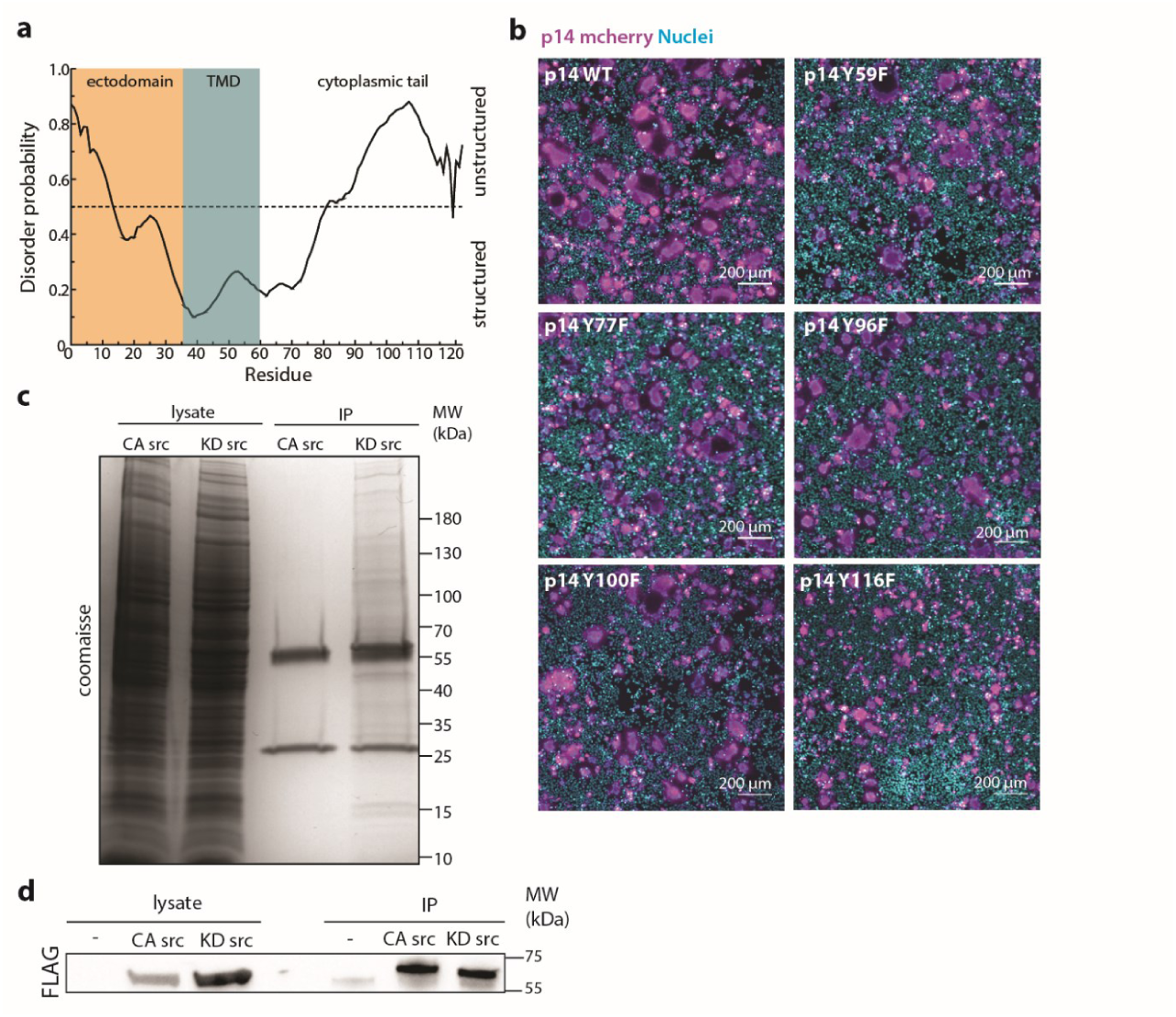
Characterization of p14 cytoplasmic tail and Y116. (a) Disorder probability of p14 cytoplasmic tail as calculated with DisEMBL (http://dis.embl.de/). (b) Representative field of view of HEK293T cells expressing p14 WT and p14 mutants with p14 labeled with C-terminus mcherry (magenta), nuclei stained with Hoechst 33342 (cyan). (c) Coomassie stain of immunoprecipitation of FLAG-tagged constitutively active src kinase (CA src) and kinase dead src kinase (KD src) from HEK293T cells. (d) Western blot of immunoprecipitation of FLAG-tagged constitutively active src kinase (CA src) and kinase dead src kinase (KD src) probed with α-FLAG.

**Figure 3 – figure supplement 1.**
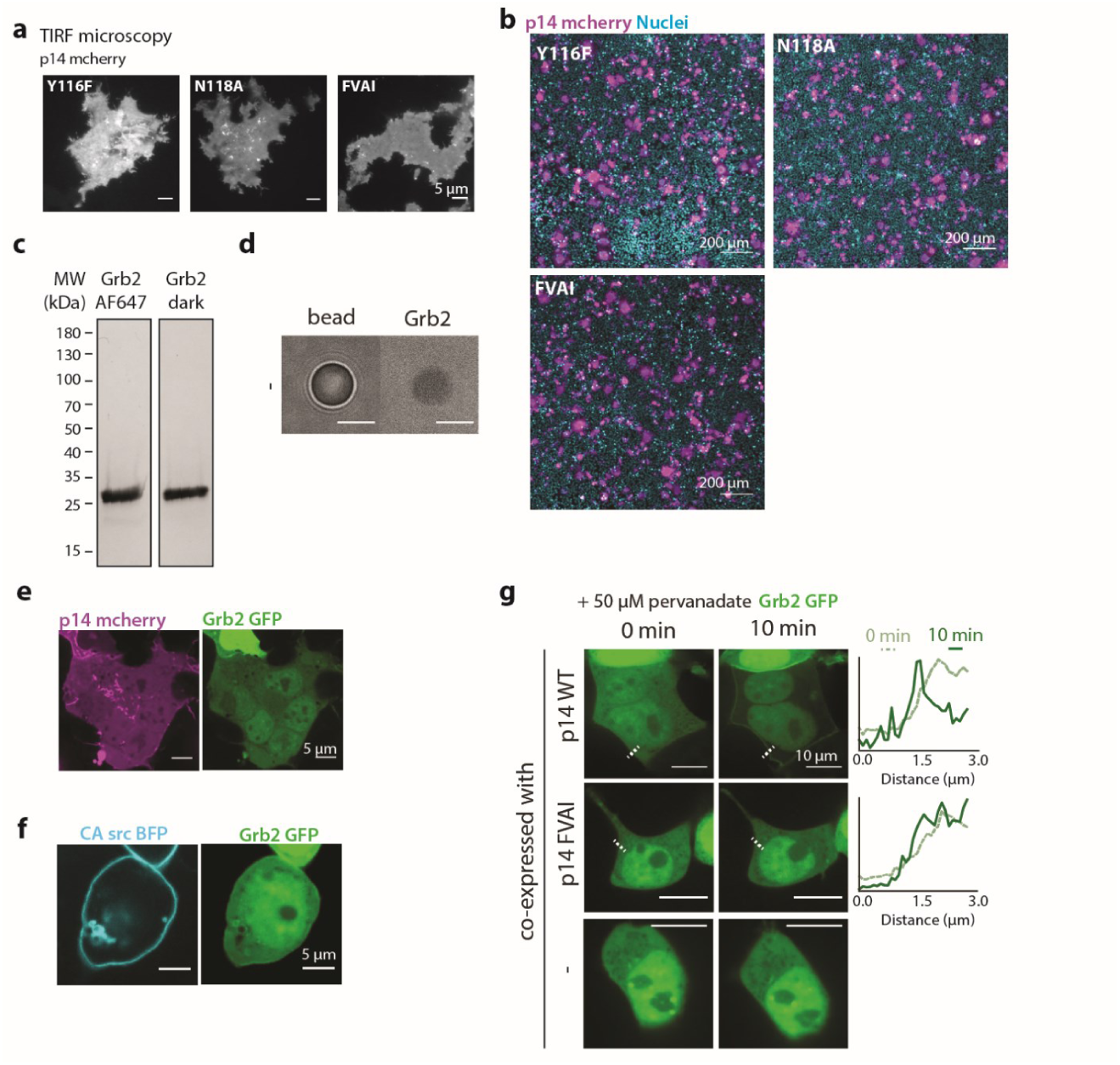
Characterization of Grb2 binding to p14. (a) p14 mutants, Y116F, N118A, FVAI, are trafficked to the plasma membrane as visualized with TIRF microscopy. (b) Representative field of view of HEK293T cells expressing p14 WT and p14 mutants with p14 labeled with C-terminus mcherry (magenta), nuclei stained with Hoechst 33342 (cyan). (c) Coomaisse stain of purified human Grb2 (right) and labeled with AlexaFluor647 (left). (d) Biotin bead incubated with purified Grb2 has minimal binding. (e) p14-WT-mcherry (magenta) co-expressed with Grb2-GFP (green) in non-treated WT HEK293T cells. (f) HEK293T cell co-expressing constitutively active src BFP (cyan) and Grb2-GFP (green) with minimal Grb2-GFP re-localized to the plasma membrane. (g) Confocal images of Grb2 enrichment to the plasma membrane of cells upon treating with pervanadate with a line scan of fluorescence intensity of each protein along the indicated white line. Grb2 does not re-localize to plasma membrane for p14 FVAI and HEK293T WT cells (bottom two rows).

**Figure 4 – figure supplement 1.**
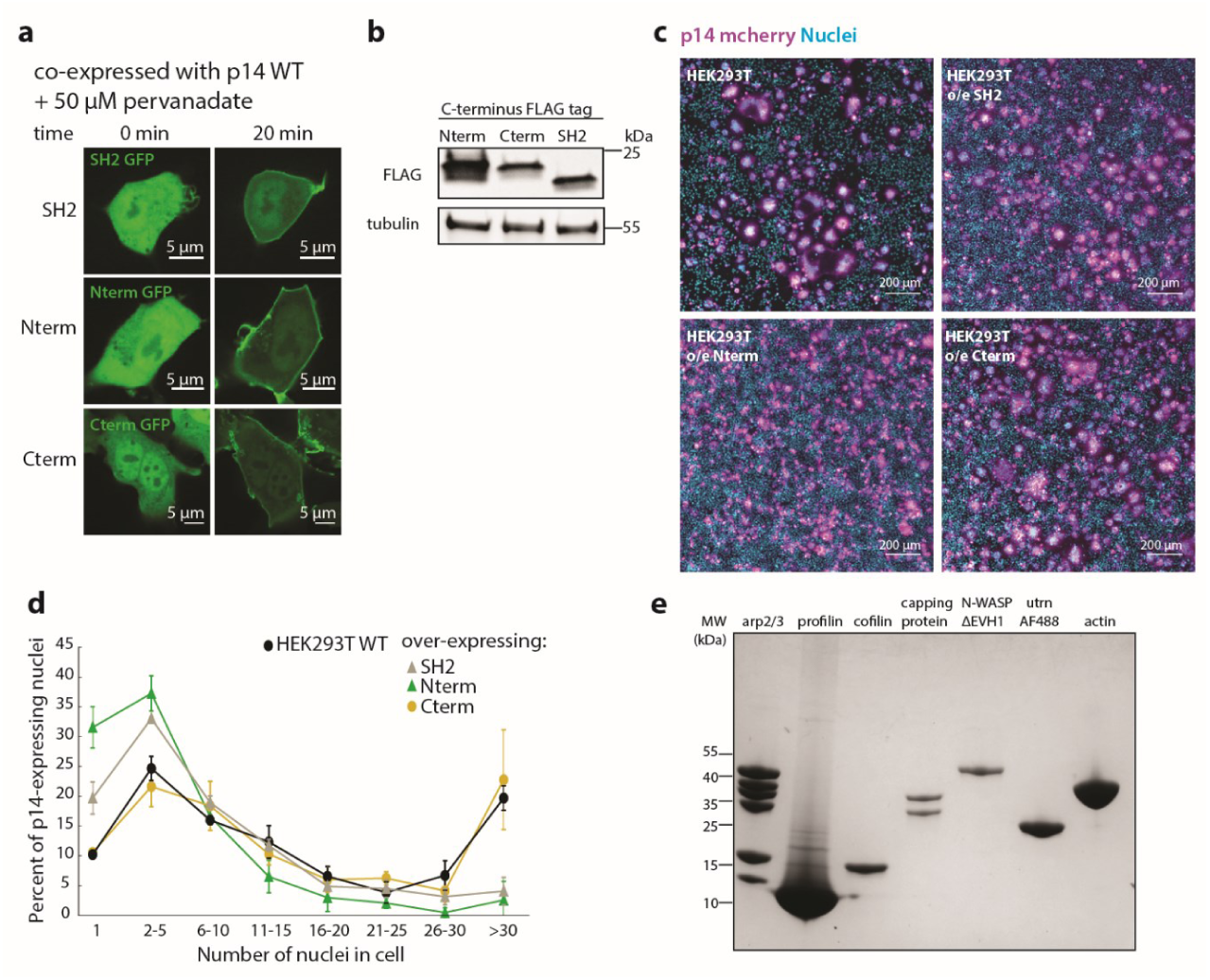
Characterization of over-expression of Grb2 mutants and purified components of actin motility assay. (a) Representative confocal images of HEK293T cells over-expressing Grb2 mutants tagged with GFP before and after addition of pervanadate to increase p14 phosphorylation. Grb2 mutants are functional, and colocalizes with p14 at the plasma membrane. (b) Western blot of HEK293T cells over-expressing FLAG-tagged Grb2 mutants. (c) Representative field of view of HEK293T cells and HEK293T cells over-expressing Grb2 mutants transfected with p14 WT with p14 labeled with C-terminus mcherry (magenta), nuclei stained with Hoechst 33342 (cyan). (d) Average nuclei count of multinucleated HEK293T cells and HEK293T overexpressing SH2 domain, N-terminus SH3 and SH2 and C-terminus SH3-SH2 domains of Grb2. p values are two-tailed, two-sample Student’s t-test to HEK293T WT cells where *p< 0.01, and **p<0.001. Error bars represent standard deviation of 3 independent transfections. (e) Coomaisse stain of SDS-PAGE of each protein used in *in vitro* actin motility assay.

**Figure 5 – figure supplement 1.**
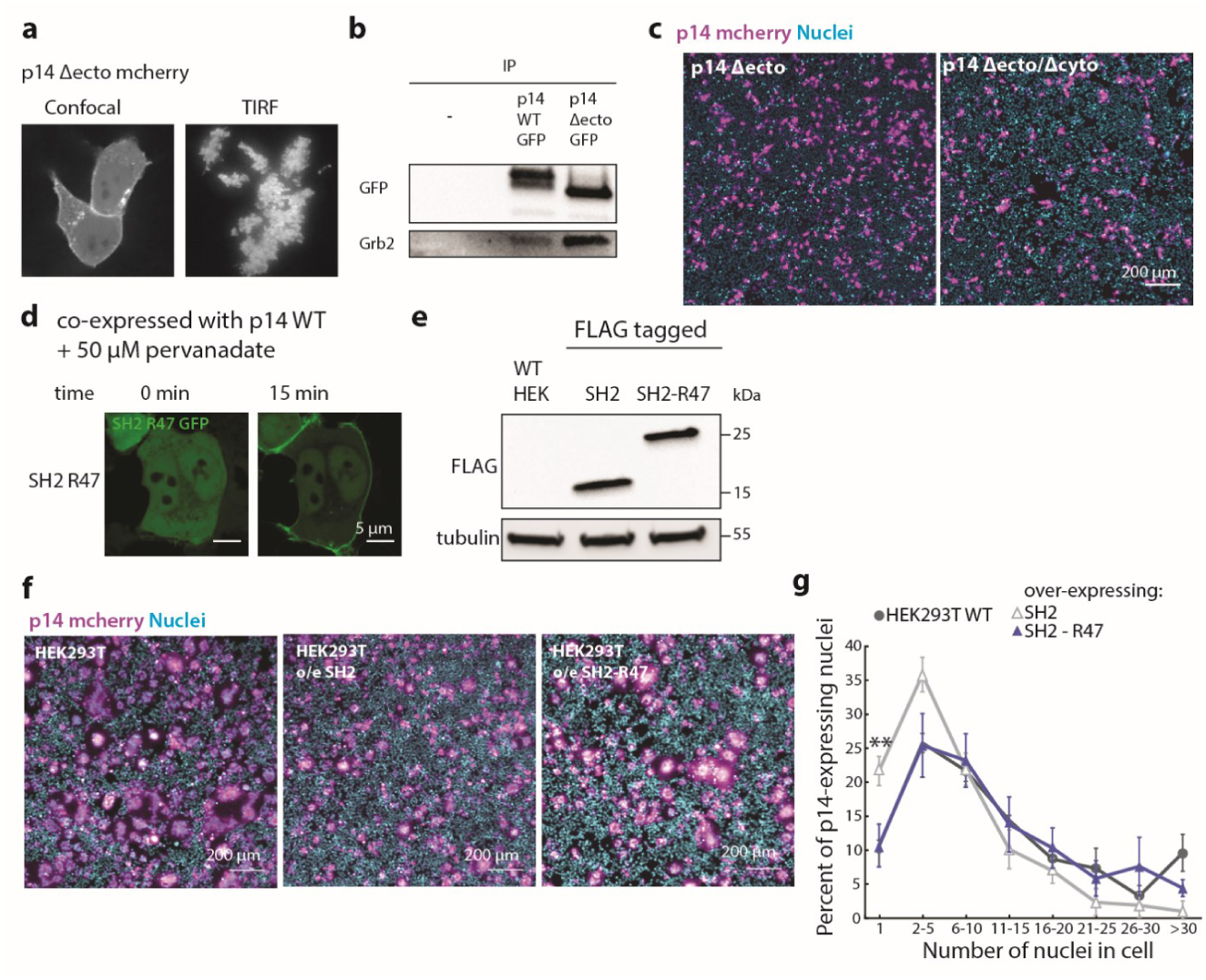
Characterization of p14 Δectodomain and direct coupling of p14 to actin assembly. (a) Representative confocal and TIRF images of p14 Δectodomain showing plasma membrane localization. (b) Western blot of co-immunoprecipitation of p14 Δectodomain and p14 WT with Grb2. (c) Representative field of view of HEK293T cells expressing p14 WT and p14 truncation mutants with p14 labeled with C-terminus mcherry (magenta), nuclei stained with Hoechst 33342 (cyan). (d) Representative confocal images of HEK293T cells over-expressing SH2-47 residues from EpsF(U) tagged with GFP and p14 WT mcherry (not shown) before and after addition of pervanadate to phosphorylate p14. (e) Western blot of HEK293T cells over-expressing FLAG-tagged SH2 and SH2-47 residues from EpsF(U). (f) Representative field of view of HEK293T cells and HEK293T cells over-expressing SH2 and SH2-47 residues from EpsF(U) transfected with p14 WT with p14 labeled with C-terminus mcherry (magenta), nuclei stained with Hoechst 33342 (cyan). (g) Average nuclei count of multinucleated HEK293T cells and HEK293T overexpressing SH2 domain and SH2-R47. p values are two-tailed, two-sample Student’s t-test to HEK293T WT cells where *p< 0.01, and **p<0.001. Error bars represent standard deviation of 3 independent transfections.

**Video 1**

Phase contrast timelapse of HEK293T cells expressing p14 WT showing extensive syncytium formation.

**Video 2**

Confocal timelapse of a HEK293T expressing p14-mcherry (magenta) fusing with a WT HEK293T cell that appears dark. Plasma membrane is marked with gpi-anchored pHluorin (green).

**Figure 1 – source data 1**

Excel Spreadsheet of counts and distribution for p14-expressing nuclei at 12 hour and 24 hour post transfection for Figure 1c.

**Figure 4 – source data 1**

Excel Spreadsheet of counts and distribution for p14-expressing nuclei of HEK293T cells over-expressing Grb2 mutants for Figure 4c.

**Figure 5 – source data 1**

Excel Spreadsheet of counts and distribution for p14-expressing nuclei of HEK293T cells over-expressing R47 constructs for Figure 5c.

